# Critical role of the periplasm in copper homeostasis in Gram-negative bacteria

**DOI:** 10.1101/2020.08.17.251918

**Authors:** Jun-ichi Ishihara, Tomohiro Mekubo, Chikako Kusaka, Suguru Kondo, Hirofumi Aiba, Shu Ishikawa, Naotake Ogasawara, Taku Oshima, Hiroki Takahashi

## Abstract

Copper is essential for life, but is toxic in excess; that is, cells must keep an optimal internal copper concentration. Under aerobic conditions, less toxic Cu(II) taken up by bacterial cells is reduced to more toxic Cu(I) in the cytoplasm. Copper homeostasis is achieved in the cytoplasm and the periplasm as a unique feature of Gram-negative bacteria. The copper efflux pumps, CopA and CusCBA export Cu(I) from the cytoplasm or the periplasm to outside of the cells in *Escherichia coli*. In addition, the periplasmic proteins, such as a multi-copper oxidase CueO, play a role in the periplasmic detoxification. While the efflux pumps are highly conserved in Gram-negative bacteria, the periplasmic proteins are diversified, indicating that copper homeostasis in the periplasm could contribute to adaptation to various living environments. However, the role of the periplasm and periplasmic proteins in regard to whole-cell copper homeostasis remains unknown. In this study, we addressed the role of the periplasm and periplasmic proteins in copper homeostasis to adapt to various ecological niches. We have used a systems approach, alternating rounds of experiments and models, to further elucidate the dynamics of copper efflux system. We measured the response to copper of the main specific copper export systems in the wild type *E. coli* strain, and a series of deletion mutant strains. We interpreted these data using a detailed mathematical model and Bayesian model fitting routines, and verified copper homeostasis. Compared with the simulation and the growth in response to copper, we found that the growth was associated with copper abundance in the periplasm. In particular, CueO unique to Gram-negative bacteria contributes both to protection against Cu(I) toxicity and to incorporating copper into the periplasmic components/proteins, resulting in maximizing the growth. These results suggest that Gram-negative bacteria have evolved to utilize the periplasm as a sensor and store for copper, in order to enable Gram-negative bacteria to adapt to a wide range of environmental copper concentrations.

## Introduction

Metals such as copper and zinc are essential for life, but toxic in excess. Metal cofactors are a prerequisite for central bioenergetic processes such as respiration, photosynthesis and nitrogen fixation (Merchant and Helmann, 2012). Metals can be bioavailable but not synthesized, and therefore complex mechanisms are required to stably maintain the concentrations of essential metals in the cell. The homeostasis of essential metals, such as copper and zinc, is achieved in a metal-specific manner in *Escherichia coli* (Chandrangsu et al., 2017; Pal et al., 2017). Zinc is imported by a high-affinity ABC-type zinc uptake system ZnuABC (Patzer and Hantke, 1998; Patzer and Hantke, 2000; Yatsunyk et al., 2008) and exported by a P-type ATPase ZntA (Rensing et al., 1997), respectively. In contrast with zinc homeostasis, copper homeostasis is achieved by the export system. In addition, copper adopts distinct redox states, either less toxic Cu(II) or toxic Cu(I). *E. coli* can survive in the digestive tract of warm-blooded animals such as the stomach and duodenum, where the concentration of Cu(I) can be high, probably less than 10 µM (Rensing and Grass, 2003).

Copper can enter *E. coli* cell without a specific import system (Rensing and Grass, 2003), and can diffuse from the periplasm to the cytoplasm without any known specific transporters (Beswick et al., 1976; Outten et al., 2001). The reducing environment of the cytoplasm leads to reduction of Cu(II) (Outten et al. 2001), and Cu(II) reductase, such as NDH-2, could reduce Cu(II) during uptake as was described in eukaryotes (Rapisarda et al., 1999; Rapisarda et al., 2002). Subsequently, it is conceivable that less toxic Cu(II) is reduced to toxic Cu(I) in the cytoplasm under aerobic conditions (Rensing and Grass, 2003). Cu(I) in the cytoplasm is regulated at most one atom per cell, because the MerR homologue protein CueR with zeptomolar sensitivity to Cu(I) functions as a Cu(I) sensor (Changela et al., 2003). The periplasm is a unique cellular compartment in Gram-negative bacteria, which is surrounded by the inner and outer membranes and has a distinctive function in substrate oxidation. The periplasm is involved in several functions such as protein oxidation and metal transport (Miller and Salama, 2018). Besides Cu(II), copper ions can be as the free Cu(I) or the binding form with proteins such as metallochaperone CusF in the periplasm (Rensing and Grass, 2003).

To achieve copper homeostasis in both the cytoplasm and the periplasm, *E. coli* carries two export systems: the Cu-efflux (*cue*) system, which transports copper from the cytoplasm to the periplasm; and the Cu-sensing (*cus*) system, which transports copper from the cytoplasm and the periplasm to the outside of cells (Rensing and Grass, 2003). In addition to the export systems, it is well known that a periplasmic multi-copper oxidase CueO comprised of the *cue* system, which oxidizes Cu(I) to Cu(II), safeguards against Cu(I) toxicity in the periplasm (Grass et al., 2001; Singh et al., 2004). The *cue* system is a primary mechanism responsible for copper resistance under both aerobic and anaerobic conditions (Rensing and Grass, 2003; Bondarczuk and Piotrowska-Seget, 2013), and the *cus* system is induced under anaerobic or copper-replete conditions (Outten et al., 2001). The *cue* system is comprised of CopA, an inner membrane Cu(I)-translocating P-type ATPase, and CueO (Petersen and Møller, 2000; Rensing et al., 2000; Grass et al., 2001; Singh et al., 2004). Expression of *copA* and *cueO* regulated by CueR is induced by Cu(I) levels in the cytoplasm (Outten et al., 2000; Stoyanov et al., 2001; Petersen and Møller, 2000). Upon exposure to copper stress such as 10 µM copper, the *cue* system is induced in order not to accumulate an excess of Cu(I); Cu(I) in the cytoplasm is transported to the periplasm by CopA, followed by the oxidization of Cu(I) by CueO. Under anaerobic or copper-replete conditions such as 500 µM copper, the *cus* system is activated; a copper-responsive two-component system CusRS induces the expression of *cusCFBA* (Yamamoto and Ishihama, 2005), encoding a multiunit copper efflux pump that spans from the outer membrane to the periplasm, which exports copper in the cytoplasm and the periplasm to the extracellular environment (Munson et al., 2000; Outten et al., 2001; Oshima et al., 2002; Franke et al., 2003). CusS senses Cu(I) in the periplasm (Outten et al., 2001), followed by the phosphorylation of CusR (Yamamoto et al., 2005). Phosphorylated CusR induces the expression of *cusC* (Rensing and Grass, 2003; Loftin et al., 2005; Kittleson et al., 2006).

Although the biochemical activity of CueO was corroborated, the effect of Cu(I) detoxification by CueO on copper stress has remained elusive. Outten et al. (2001) reported that while the expression of *cueO* was induced at a copper concentration of 10 µM, the disruption of *cueO* had little effect on the growth compared with the growth in the Δ*copA* strain even upon exposure to a higher copper concentration of 500 µM (Outten et al., 2001). Hernández-Montes et al. (2012) reported that the periplasmic proteins, such as CueO, involved in copper homeostasis are extensively diversified among 268 gamma proteobacteria, while copper transport from the cytoplasm to the periplasm by CopA and from the cytoplasm and the periplasm to the outside of the cells by CusCBA is highly conserved (Hernandez-Montes et al., 2012). Although this finding indicates that periplasmic proteins function in a species-specific manner, the role of the periplasm in copper stress is also unknown. These raise the question of whether Cu(I) detoxification by CueO serves as protection against copper stress, and whether there are the common features of periplasmic components/proteins in copper homeostasis.

In this study, we addressed the role of CueO and the periplasm in sensing and storing of copper. We initially performed ChIP-chip analysis to confirm the regulons controlled directly by CueR and CusR in *E. coli*, and measured the transcription responses to copper of the copper efflux systems by Lux reporter analysis. We then derived a mathematical model, and integrated the model with both published data and our current experimental data using a Monte Carlo Markov chain approach to evaluate model fits and infer the plausible ranges of parameter values. Compared the stochastic simulations with the growth measurements, we suggested that CueO unique to Gram-negative bacteria contributes both to protection against Cu(I) toxicity and to incorporating copper into the periplasmic components/proteins. Taken together with comparative genome analysis, we propose that Gram-negative bacteria have evolved to utilize the periplasm as a sensor and store for copper, contributing to the expansion of living niche with a wide range of environmental copper concentrations.

## Results

### Confirmation of the genes directly regulated by CueR and CusR

In order to identify a set of genes directly regulated by CueR and CusR of the *cue* and *cus* systems, respectively, we performed ChIP-chip analysis. The *E. coli* TM01 and TM02 strains expressing 3x Flag-tagged CueR and CusR, respectively, were grown aerobically in Defined Medium A (DMA) supplemented with 0 µM, 750 µM and 2 mM CuSO4. Seeking the binding peaks of CueR and CusR, we found that two CueR-binding peaks were located at the upstream regions of *copA* and *cueO* across the entire *E. coli* chromosome at 750 µM and 2 mM CuSO4 (Figure 1B). We found that one CusR-binding peak was located at an intergenic and upstream region of *cusCFBA* and *cusRS* operons at 2 mM CuSO4 (Figure 1B). This result was consistent with a previous finding that the CusR binding sequence (CusR box) was located only at that region (Yamamoto et al., 2005).

**Figure 1.**
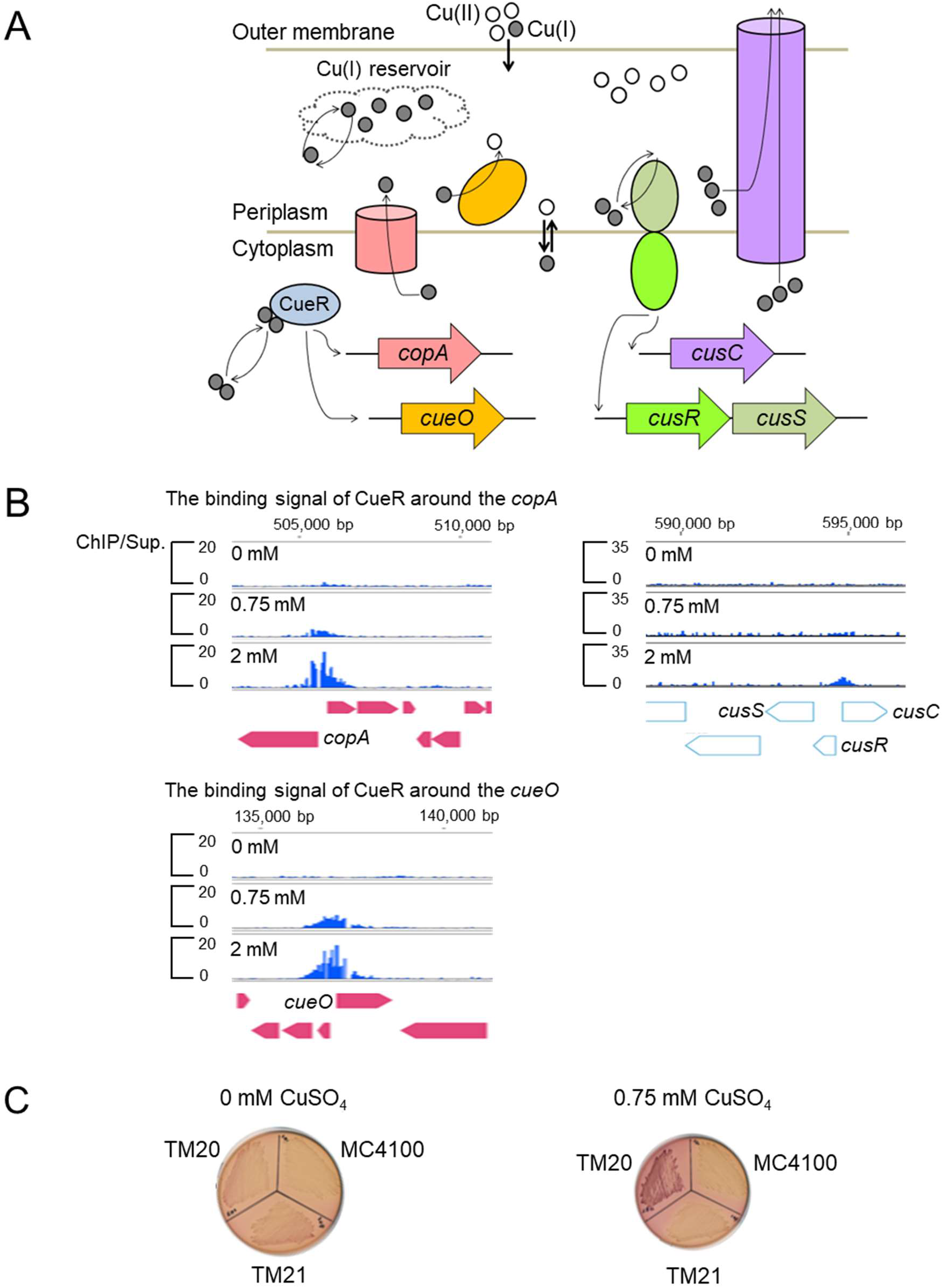
(A) Schematic figure of copper efflux system, i.e. *cue* and *cus* systems in *E. coli*. Cu(II); white circle, Cu(I); black circle, CueR; aqua ellipse, CopA; pink cylinder, CueO; yellow ellipse, CusC; purple cylinder, CusS; light green ellipse, and CusR; yellow-green ellipse. (B) CueR and CusR binding sites on the genome in DMA^0^, DMA^750^, and DMA^2000^ identified by ChIP-chip. The values on the vertical axes indicate the binding magnitudes of ChIP DNA relative to those of supernatant DNA, and the horizontal axes indicate the genomic coordinate. (C) Expression of *cusR-lacZ* by β-galactosidase activity. MC4000, TM20 (MC4100 *cusR*-*lacZ* fusion) and TM21 (MC4100 *cusRS*::*km^r^ cusR*-*lacZ* fusion) were streaked on MacConkey agar plates supplemented without (left) and with 750 µM CuSO4 (right).

Since auto-regulation of CusR remains unclear, we investigated the auto-regulation of CusR by the plate assay of *cusR-lacZ* fusion strains. The TM20 strain formed red colonies on MacConkey agar plates supplemented with 750 µM CuSO4, while the MC4100 and TM21 strains formed white colonies (Figure 1C). In contrast, we confirmed that all strains formed white colonies on MacConkey agar plates without CuSO4. Our results clearly indicated that CusR directly and positively regulated *cusRS* in the presence of CuSO4. Taken together, we concluded that four promoters, that is, *copA, cueO, cusCFBA,* and *cusRS* are regulated by CueR and CusS, of the *cue* and *cus* systems in *E. coli*.

### Time course of promoter activities following addition of copper

To elucidate the transcription responses of the *cue* and *cus* systems, the promoter activities of three genes, *copA* (henceforth called P*copA*), *cueO* (P*cueO*), and *cusCFBA* (P*cusC*) in the wild-type MG1655 (WT) strain, and its derivatives Δ*copA*, Δ*cueO*, and Δ*cusC* strains, were measured using a Lux reporter system (Burton et al., 2010; Takahashi et al., 2015). Excess external copper conditions included the addition of either 100 µM, 200 µM, 300 µM, or 400 µM CuSO4 to DMA (henceforce called DMA^100^, DMA^200^, DMA^300^, or DMA^400^). We confirmed no severe growth defects in all strains (data not shown).

Figure 2J shows the maximum induction levels of promoters relative to the promoter signal at 0 min in the WT strain. The maximum induction levels of P*cusC* were associated with the addition of CuSO4 in a concentration-dependent manner. In contrast, we observed little change in maximum induction levels of P*copA* above DMA^300^, and P*cueO* above DMA^200^, respectively. These results suggested that the *cue* system could sense the differences between lower external copper concentrations such as DMA^100^ and DMA^200^, while the *cus* system could sense the differences between higher external copper concentrations such as DMA^300^ and DMA^400^, which could not be distinguishable by the *cue* system.

**Figure 2.**
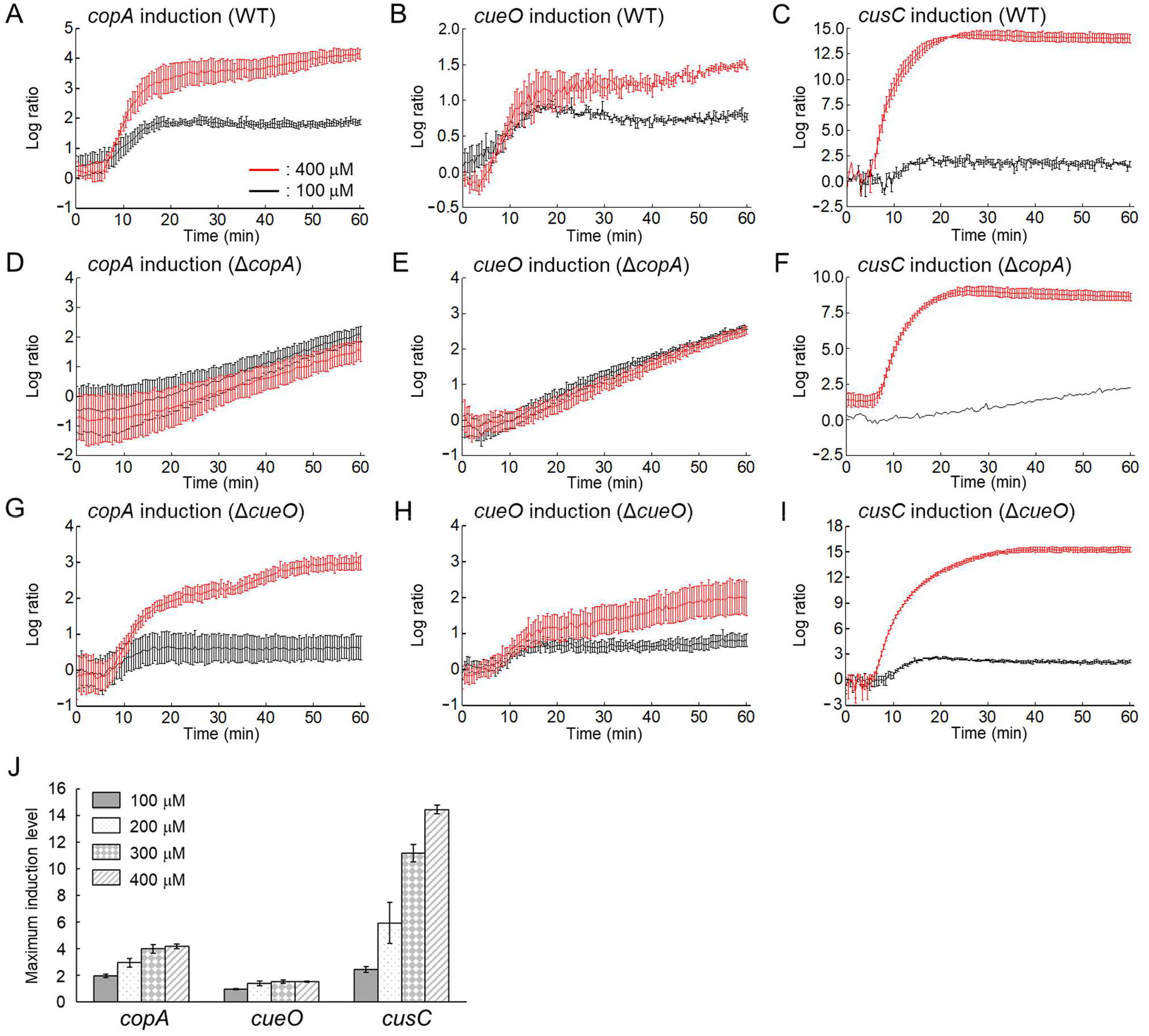
(A–I) Time series of P*copA*, P*cueO*, and P*cusC* following addition of 100 µM (black lines) and 400 µM (red lines) copper (log to base 2 ratios to the value at *t*=0). The remaining experimental data are shown in Figure S1. Error bars indicate s.d. (*n*=3). (J) Maximum induction levels of P*copA*, P*cueO*, and P*cusC* relative to the data at 0 min in the WT strain (log to base 2 ratios to the value at *t*=0).

Next we investigated the effects of a CopA deletion on the inductions of P*copA*, P*cueO*, and P*cusC*. All tested genes were affected by the CopA defect (Figure 2D–F). CopA is the only known chromosomally encoded Cu(I) efflux pump from the cytoplasm to the periplasm in *E. coli*. The inductions of P*copA* and P*cueO* continued for 60 min after addition of CuSO4 in the Δ*copA* strain, although the inductions of these promoters in the WT strain plateaued after 20 min (Figures 2A, B, D and E). This could be because the deletion of CopA causes Cu(I) accumulation in the cytoplasm, resulting in the continued activation of P*copA* and P*cueO* in DMA^100^ and DMA^400^. We also expected little change of P*cusC* induction, because the expression of *cusC* was positively regulated by the two-component system CusRS which was activated through the increase of Cu(I) in the periplasm (Rensing and Grass, 2003). Interestingly, P*cusC* was induced in the Δ*copA* strain following addition of CuSO4 (Figure 2F), supporting the activity of alternative Cu(I) exporters such as AcrD and MdtABC from the cytoplasm to the periplasm in addition to CopA (Nishino et al., 2010; Hobman and Crossman, 2015). The induction levels of P*cusC* were, however, drastically reduced compared to those in the WT strain (Figures 2C and F).

In addition, the effects of a CueO deletion on the induction of P*copA*, P*cueO*, and P*cusC* were investigated (Figures 2G–I). We hypothesized that as CueO plays an important role in oxidation of Cu(I) to Cu(II) in the periplasm, the disruption of CueO had no impact on the inductions of P*copA* and P*cueO*. As expected, we observed similar expression profiles of P*copA* and P*cueO* in both the WT and Δ*cueO* strains. In contrast to the effects of the CueO defect on P*copA* and P*cueO*, it was thought that the induction of P*cusC* could occur at lower concentration of CuSO4 in the Δ*cueO* strain compared to the WT strain, because deletion of *cueO* causes a decrease in periplasmic Cu(I) oxidation activity, leading to higher Cu(I) levels in the periplasm. As expected, P*cusC* in the Δ*cueO* strain exhibited 4.4-fold change relative to that in the WT strain in DMA^100^.

Finally, we investigated the effects of a CusC deletion on the induction of P*copA*, P*cueO*, and P*cusC* (Figures S1A–C). Copper concentrations we tested were lower than the concentration for optimum growth, that is, 750 µM (Kershaw et al., 2005). As expected, the disruption of CusC had no effect on the maximum induction levels of P*cusC* relative to the promoter signal at 0 min (Figures 2J and S1D). Consistent with previous observation (Munson et al., 2000), the *cus* system was induced at relatively high concentrations of CuSO4, while the *cue* system responded to relatively low concentration of CuSO4 under aerobic conditions.

### A new mathematical model for *in vivo* copper homeostasis

In order to understand the dynamics of the *cue* and *cus* systems, in conjunction with published data, we developed a mathematical model that describes the molecular process of copper homeostasis including Cu(I) concentrations in the cytoplasm and the periplasm (Figure 1A). In the model, we mainly focused on the interactions between the *cue* and *cus* systems which dominate the dynamics of Cu(I) in the cell. We set three variables for expressing copper concentrations as Cu(I) in the cytoplasm, Cu(I) in the periplasm, and Cu(II) in the periplasm, leading us to incorporate the spatiotemporal dynamics of Cu(I) and Cu(II) in the cell. Furthermore, we use the model to test the role of CueO in copper homeostasis. In the model, we assumed two features. First, that there is a Cu(I) reservoir in the periplasm that protects the cells against Cu(I) toxicity. This is because there exist proteins acting as the copper reservoir, such as CusF, which play a role in keeping vast majority of Cu(I) in the periplasm to transfer Cu(I) to other proteins (Loftin et al., 2005, Kittleson et al., 2006, Pal et al., 2017). Second, we hypothesize that CusS phosphorylates and dephosphorylates its two-component partner CusR in the same manner as the other sensor kinases (Miyashiro and Goulian, 2008, Ray and Igoshin, 2010), i.e. CusS acts as a bifunctional kinase (Figure S2 and equations 11–16 in Materials and Methods). Consequently, we derived the mathematical model including 16 variables (see Materials and Methods).

### Model fitting to data

Model fitting to data including our Lux data and other published data has been accomplished using the Metropolis-Hastings algorithm for parameter inference. Convergence plots from the simulations and posterior distributions for all parameters are shown in Figure S3. Point estimates and ranges for all parameter values are shown in Table S1. Independently of copper concentrations, P*copA* and P*cueO* activities exhibited the linear increase in the Δ*copA* strain (Figures 2D, 2E, S1A, and S1B), indicating that CueR could be continuously activated by accumulated copper in the cytoplasm. We thought that deletion of CopA caused Cu(I) accumulation in the cytoplasm due to lack of cytoplasmic copper export, and P*copA* and P*cueO* inductions occurred before addition of CuSO4 (Figure S1E). These data were excluded from the parameter inference to focus on the concentration-dependent regulations.

### The model fits a zeptomolar sensitivity of the transcriptional factor *in vitro* data

The model fits very closely (Figure 3A) to the published zeptomolar sensitivity of CueR for Cu(I) (Changela et al., 2003), providing confidence in the model results, especially given the high sensitivity of the molecular switch. The binding of two Cu(I) ions is required to convert inactive CueR to active CueR at rate *l_1_*; the median value was 0.136 nM^−2^s^−1^. The active form reverts to the inactive form at rate *l_2_*; the median value was 4.06×10^−25^ s^−1^ (Table S1 and equation 4 in Materials and Methods). Consequently, the half-maximal induction of CueR was estimated at 1.72×10^−21^ M, consistent with the reported data, 2×10^−21^ M (Changela et al. 2003).

**Figure 3.**
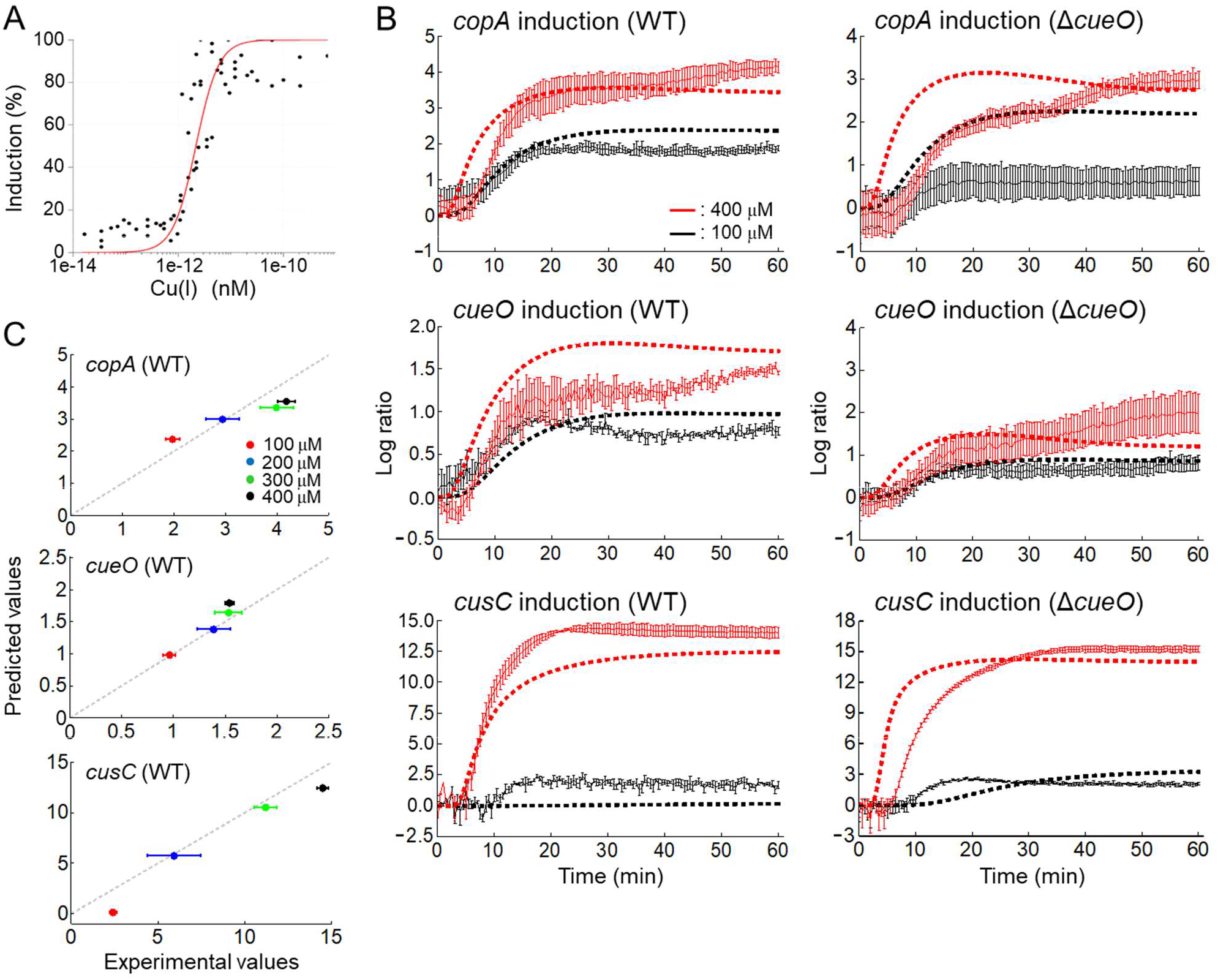
(A) Model fit to CueR induction data from Changela et al, 2003. Black dot is the experimental value, and red line is the model value. (B) Time series of induction of P*copA*, P*cueO*, and P*cusC* following addition of 100 µM copper (black lines) and 400 µM copper (red lines) (log to base 2 ratios). The solid line indicates the experimental data (error bar is s.d.), which are shown in Figure 2. The dotted line indicates the theoretical data with the best fit parameters. The remaining model fits are shown in Figure S4. (C) Experimental and theoretical maximum induction levels of P*copA*, P*cueO*, and P*cusC* in the WT strain (log to base 2 ratios to the value at *t*=0).

### The model fits the P*copA*, P*cueO*, and P*cusC*

The model replicated experimental data for *in vivo* inductions of P*copA*, P*cueO* and P*cusC* in all strains (Figures 3B and S4), with a very high goodness of fit (*R^2^* = 88.2%) associated with the promoter activities in all strains under all conditions. There were two exceptions where the fit was less good: the model predicts higher induction of P*copA* in the Δ*cueO* strains, while the model fits to all inductions of P*copA* in the WT and Δ*cusC* strains. In addition, although the model fits to the inductions of P*cusC* in the WT and Δ*cueO* strains, the model fit of the induction of P*cusC* in the Δ*cusC* strain is less good (Figure S4C): the model predicts greater induction (15.99 [log_2_ ratio] over the time course) than the experimental data (6.78 [log_2_ ratio] over the time course). The model also gives a reasonable fit to the maximum inductions of P*copA*, P*cueO* and P*cusC* (Figure 3C). Overall, these results give considerable confidence in the model processes and parameter estimations.

### Prediction of copper abundance in the periplasm from a stochastic model

P*cusC* exhibited the highest level of induction among the genes we tested in the WT strain (Figures 3B and S1C). We observed a similar behavior of P*cusC* in the Δ*cueO* strain, indicating that Cu(I) accumulates in the periplasm following addition of copper regardless of whether CueO exists. In order to explore copper atom levels, i.e. Cu(I) and Cu(II), in the periplasm in single cells, we developed a stochastic model. The model contains exactly the same processes as the ODE model described above, but uses a discrete chemical reaction scheme to describe them (Gibson and Bruck, 2000), and thus captures intrinsic variability due to molecular events (Swain et al., 2002).

To reveal the dynamics of copper in the WT and Δ*cueO* strains, we conducted stochastic simulations following addition of a wide range of copper concentrations from 0 µM to 6,000 µM. We adopted a copper concentration of 6,000 µM as the upper limit to examine the dynamics under extremely high copper concentration, where the growth was experimentally confirmed (Kershaw et al. 2005). In the WT and Δ*cueO* strains, Cu(I) and Cu(II) abundance in the periplasm increased with the increase in the external concentrations of copper (Figures 4A and B). In contrast, the cytoplasmic free Cu(I) was at most one molecule per cell under all tested conditions, indicating that copper homeostasis in the cytoplasm could be achieved.

**Figure 4.**
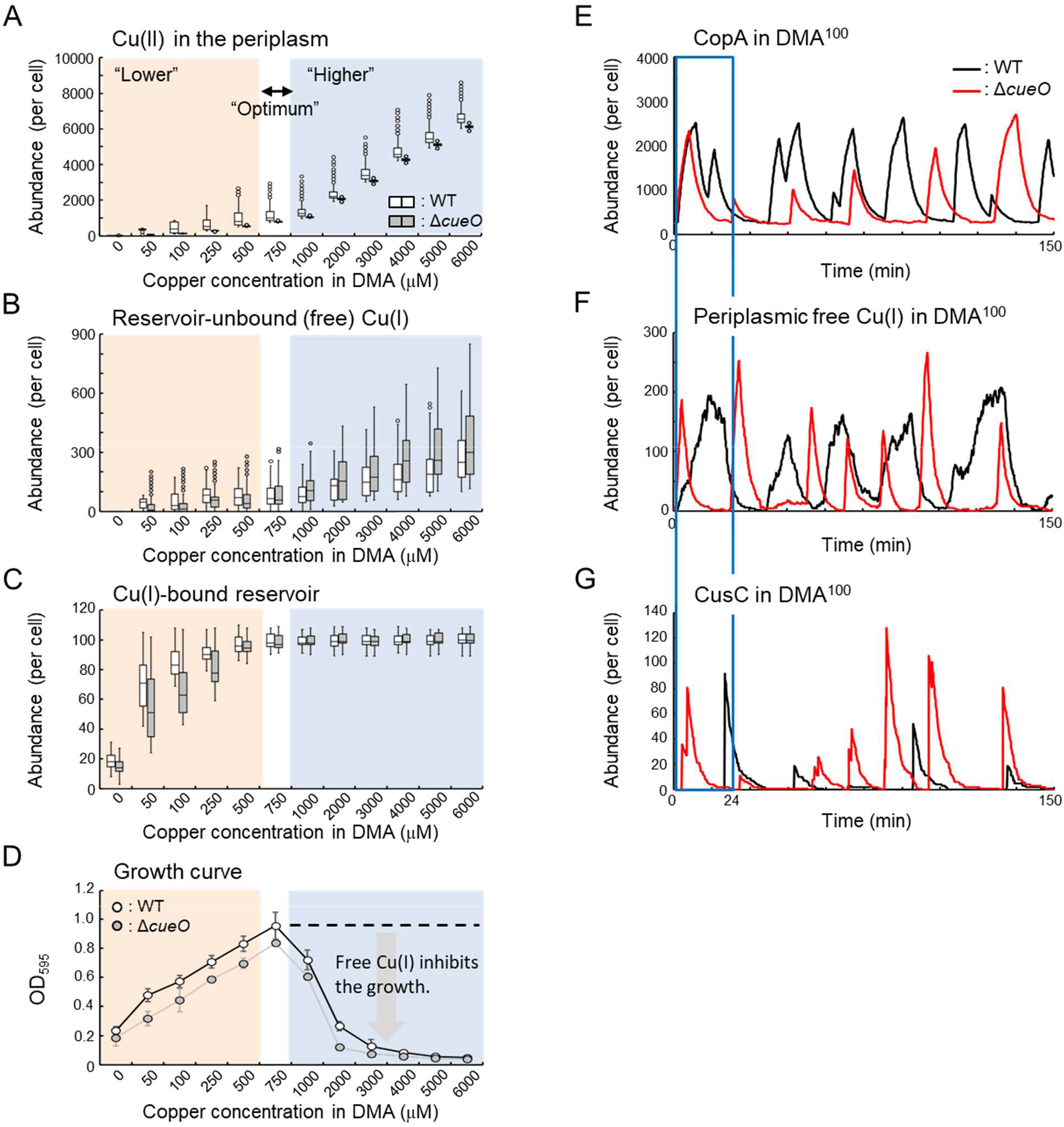
(A, B, C) Boxplots of the abundance of periplasmic free Cu(II), reservoir-bound Cu(I), and free (reservoir-unbound) Cu(I) in the WT (black) and Δ*cueO* (gray) strains. (D) Effect of the copper concentration in DMA upon the growth of WT (●) and Δ*cueO* strains (◯). The experiment was performed in triplicate, and error bars indicate s.d. (E–G) Realization of the stochastic simulation showing the abundance of CopA, periplasmic free Cu(I), and CusC in the WT (pink) and Δ*cueO* (light green) strains in DMA^100^. The profiles from 0 min to 24 min were surrounded by the red line.

### *cueO* contributes both to protection against Cu(I) toxicity and to incorporating copper in the periplasm

The abundance of Cu(II) was higher in the WT strain than in the Δ*cueO* strain (Figure 4A). This could be caused by the oxidation of Cu(I) to Cu(II) by CueO in the WT strain. Under external copper concentrations ranging from 0 to 500 µM (designated “Lower”), the abundance of Cu(I) was lower in the WT strain than that in the Δ*cueO* strain (Figure 4B). This could be because Cu(I) is exported by CusC in the Δ*cueO* strain. Looking at the profiles of CopA and periplasmic Cu(I) abundance in the Δ*cueO* strain, the induction of expression of CopA and the increase in the Cu(I) level occurred simultaneously (Figures 4E and F), followed by the expression of CusC (Figure 4G), resulting in lower Cu(I) levels in the Δ*cueO* strain. In contrast, the expression of CopA was followed by the oxidation of Cu(I) by CueO in the WT strain. Consequentially, the timing of expression of CusC was delayed (Figure 4G). Interestingly, we observed higher abundance of the Cu(I)-bound reservoir in the WT strain than that in the Δ*cueO* strain. In other words, the WT strain maintains higher levels of copper in the periplasm. This is likely due to the delay of expression of CusC caused by Cu(I) oxidation through CueO.

We assumed the existence of the reservoir molecules bound to Cu(I) in the periplasm, such as CusF, which transfers Cu(I) to other proteins, leading to incorporation of copper required for various functions. This led us to the hypothesis that the abundance of the Cu(I)-bound reservoir is linked to the growth of *E. coli*. To address the relationship between the growth and copper levels in the periplasm, we measured the growth (OD595) of WT and Δ*cueO* strains in DMA with different concentrations of copper (Figure 4D). We confirmed no differences of cell sizes between two strains (data not shown). Optimum growth of WT and Δ*cueO* strains occurred in DMA^750^ consistent with the previous report (Kershaw et al., 2005). The Δ*cueO* strain had a similar growth profile to the WT strain, although the Δ*cueO* strain exhibited poor growth compared with that of the WT strain. We found that the growth was associated with the abundance of the Cu(I)-bound reservoir (Figure S6A). The growth data for the WT strain in DMA^100^, DMA^250^, and DMA^500^ was almost identical to those of the Δ*cueO* strain in DMA^250^ DMA^500^, and DMA^750^, respectively, indicating that the growth of the WT strain was similar to that of the Δ*cueO* strain if the WT and Δ*cueO* strains accumulate a similar abundance of Cu(I) in the Cu(I)-bound reservoir.

In contrast with the “Lower” conditions, there were no differences in the abundance of Cu(I)-bound reservoir between the WT and Δ*cueO* strains under external copper concentrations over 1,000 µM (designated “Higher”) (Figure 4C). Instead, we observed a relationship between the growth and free Cu(I) abundance (Figure S6B). The model predicted the median of 149 free Cu(I) atoms per cell in the WT strain in DMA^3000^, where the growth of WT strain exhibited the OD595 value of 0.127. These values were close to those of the Δ*cueO* strain in DMA^2000^, that is, the OD595 value of 0.120 and the median of 146 free Cu(I) atoms per cell. This indicates that the strain with a particular OD595 value could be characterized by the Cu(I) abundance in the periplasm. Taken together, CueO may contribute both to protection against Cu(I) toxicity and to incorporating copper into the periplasmic components/proteins, resulting in maximizing the growth.

### Comparative genome analysis shows *cueO* and *cusC* were acquired in ***Proteobacteria***

Finally, we investigated how the transporters involved in copper and zinc homeostasis are evolutionally conserved among four bacteria in three different phyla including *Proteobacteria*, *Cyanobacteria* and *Firmicutes*, i.e. *E. coli*, *Synechococcus elongatus*, *Bacillus subtilis*, and *Staphylococcus aureus*. We identified 1,989 orthologous groups. Interestingly, no orthologous genes to *cusC* was found in *S. elongatus*, *B. subtilis*, and *S. aureus*, while *copA* is a single-copy gene among all tested species (Figure 5). *zntA* was found in *E. coli* and *B. subtilis*, while two orthologues of *znuC* in *S. elongatus*, *B. subtilis*, and *S. aureus* were classified into the same cluster. Although the orthologous gene of *cueO* was *cotA* in *B. subtilis*, CueO and CotA had 23% amino acid sequence identity. Since it is conceivable that *cotA* functions as laccase (Hullo et al. 2001), we concluded that *cueO* exists only in *E. coli*. Taken together, bacteria with periplasm could acquire *cusC* which exports Cu(I) in the periplasm and the cytoplasm to protect against Cu(I) toxicity, resulting in expansion of living niche such as copper-replete environments. Furthermore, acquirement of *cueO* makes it possible to safeguard against Cu(I) toxicity and to incorporat copper, resulting in rapid growth.

**Figure 5.**
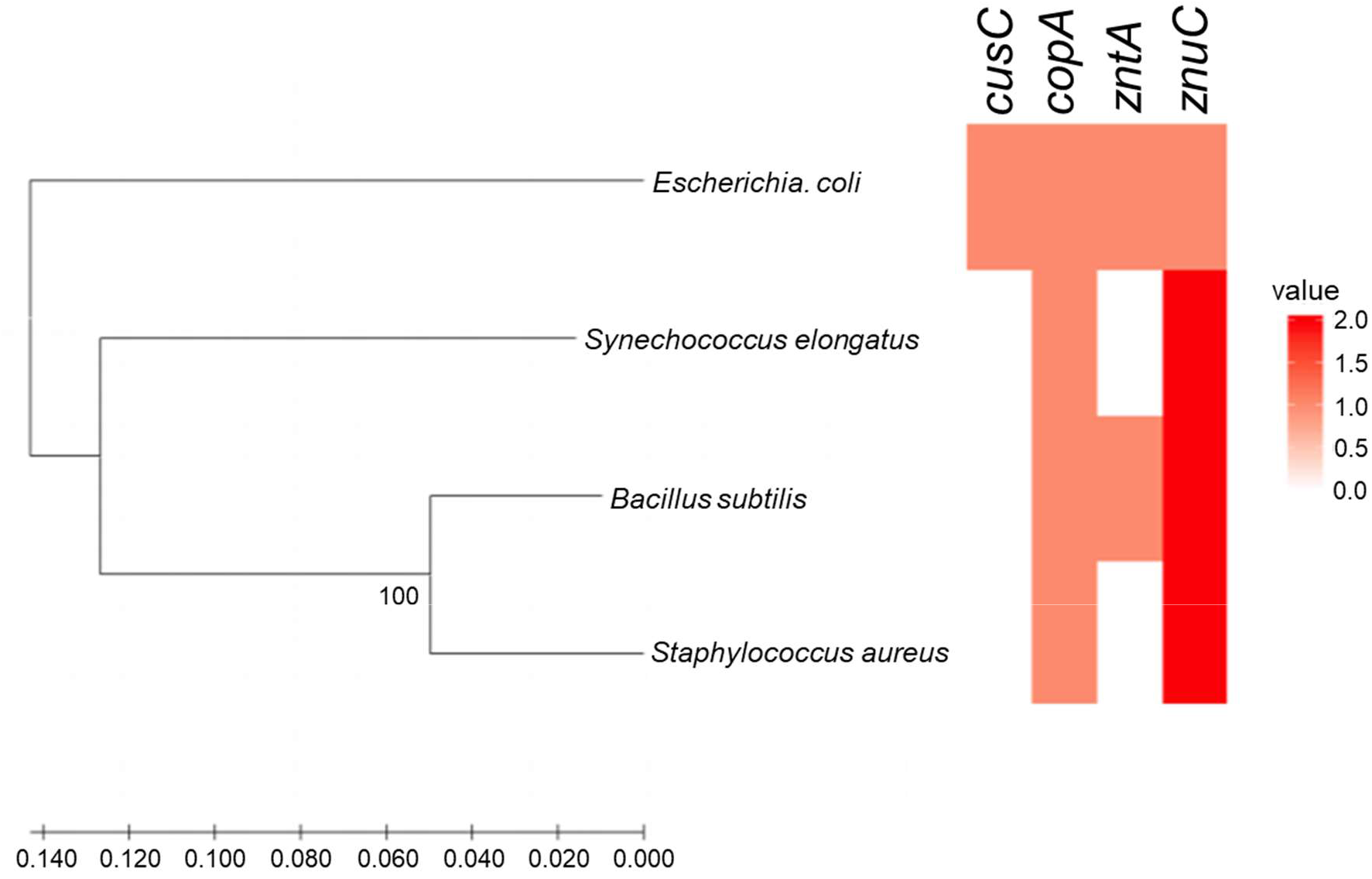
The presence of transporters involved in copper and zinc homeostasis. The phylogenetic tree was constructed based on 16S rRNA. The heatmap shows the number of genes in the orthologous groups, ranging from 0 to 2.

**Figure 6.**
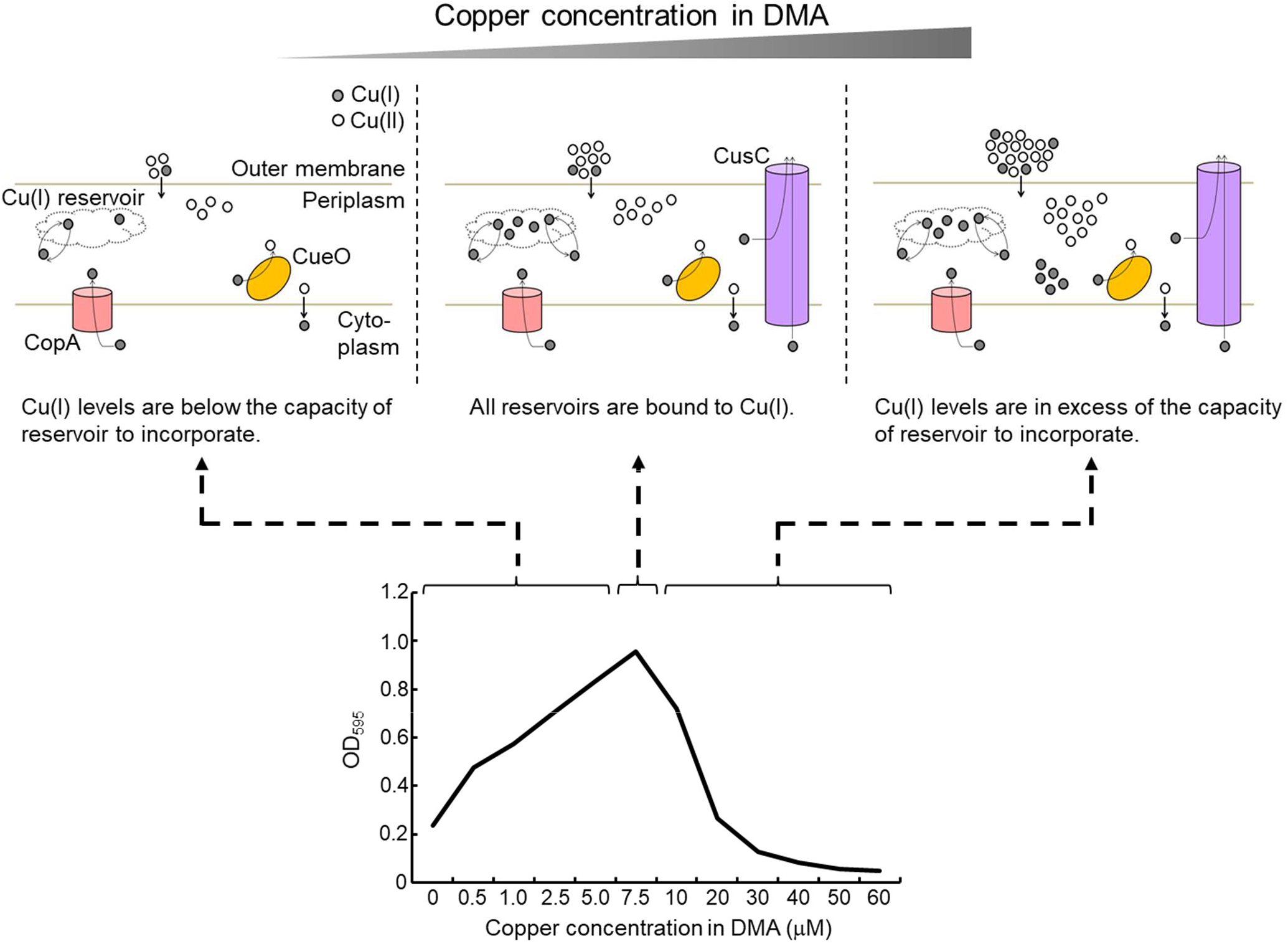
A schematic figure representing the relationship between the growth and periplasmic Cu(I) level. Under the lower copper condition, the growth is associated with the abundance of copper-bound reservoir (left). At the copper concentration of 750 µM, all reservoir molecules are bound to Cu(I), resulting in the optimum growth (middle). Under the higher copper condition, Cu(I) becomes toxic, resulting in the inhibition of growth (right).

## Discussion

We aimed to develop a detailed understanding of the responses of copper homeostasis in *E. coli* through a Systems Biology approach. Copper homeostasis is achieved through two export systems, namely, the *cue* and *cus* systems. First, to derive a mathematical model, we confirmed four operons, that is, *copA*, *cueO*, *cusCFBA*, and *cusRS* are regulated by the *cue* and *cus* systems by ChIP-chip experiments. Second, we generated novel experimental data on four strains, including time course transcriptional responses of copper efflux genes following addition of copper to the growth media in the wild type *E. coli* strain and a series of deletion mutant strains. These data, along with literature data, have been integrated with a detailed, mechanistic, mathematical model, using a Monte Carlo approach to fit this model to the data. Our model comprising of 16 equations fits the experimental data well; the model reproduced the zeptomolar sensitivity of CueR for Cu(I), and the model fits had an *R^2^* value of 88.2% for time course data. Furthermore, we validated our model in regard to the protein abundance. It was reported that the abundance of CusB transcribed together with CusC was approximately 144 molecules per cell by using LC-MS/MS (Ishihama et al., 2008). Concordantly, our simulation predicted the abundance of CusC ranging from 0 to 450 molecules per cell (Figure S5). Thus, we concluded that our model describes copper homeostasis in *E. coli*, in particular, the copper stress response and the number of molecules in the periplasm.

We assumed the CusS is bifunctional. For the dephosphorylation activity of CusS *m_8_*, the median value was 46.8 s^−1^, while for the kinase (phosphorylation) activity of that *m_6_*, the median value was 49.9 s^−1^ (Table S1), indicating the phosphatase and kinase activities of CusS was nearly identical. Although we observed that P*cusC* was induced after a few minutes following the inductions of P*copA* and P*cueO* (Figures 2A–I and S1A–C), the model did not reproduce the delay of P*cusC* when the parameters of kinase activity *m_6_* and dephosphorylation activity *m_8_* were set to be 49.9 s^−1^ and 0.1 s^−1^, respectively. Furthermore, the model without the assumption of CusS bifunctionality was not able to reproduce the delay of P*cusC* (data not shown). This result suggests that the bifunctional activity of CusS is important in determining the timing of *cusC* induction. Modulation of phosphatase and/or kinase activities of histidine kinase may affect the equilibrium between active (phosphorylated) and inactive (unphosphorylated) forms of the response regulator, and therefore alter the dynamics of target genes. It is reported that other histidine kinases phosphorylate and dephosphorylate their partner response regulators, for example, the histidine kinase PhoQ and its pertoner response regulator PhoP (Miyashiro and Goulian, 2008), EnvZ and OmpR (Batchelor and Goulian, 2003), and the TorS and TorR systems (Carey et al., 2018) in *E. coli*, supporting the bifunctional activity of CusS.

We introduced the reservoir molecules for the periplasmic Cu(I). This is reasonable because a small (110-aa) periplasmic protein CusF binds with high affinity to Cu(I) but not Cu(II) (Kittleson et al., 2006), and may function as a metallochaperone, bringing Cu(I) from the periplasm to other proteins (Loftin et al., 2005). That is, a possible role of CusF may be connected with the specific selection of Cu(I) to prevent its undesirable removal (Bagai et al., 2008; Loftin et al., 2009). We considered that the predicted abundance of reservoir molecules was reasonable, because 110 reservoir molecules was close to the abundance of CusC (Figure S3 and Table S1), consistent with the report that the abundance of CusF is almost the same as that of CusB (Loftin et al., 2005). Given that any molecules have not been reported for sequestering the periplasmic Cu(II), we did not assume any reservoir molecules for the periplasmic Cu(II). Assuming existence of the reservoir molecules for the periplasmic Cu(II) in our pilot model, the model fit was worse (data not shown). We considered that Cu(II) is not targetted to be sequestered because Cu(II) is less toxic to the cell.

Little is known about the role of CueO in copper homeostasis, although it is essential and the biochemical activity of CueO is deeply understood. Our simulation shows that the oxidation of Cu(I) by CueO moderates the accumulation of periplasmic Cu(I), leading to the delay of CusC induction. Thus, the abundance of Cu(I)-bound reservoir was higher in the WT strain than that in the Δ*cueO* strain (Figure 4C). Moreover, as the *cueO* disruption is expected to generate a higher level of Cu(I) in the periplasm, the induction of P*cusC* could occur at lower concentration of CuSO4 in the Δ*cueO* strain compared to the WT strain. That is, CueO may contribute to the delay of induction of CusC. This could be a mechanism for incorporating Cu(I) in the periplasm under the lower copper condition. It is suggested that CueO modulates sequestering or exporting of Cu(I) by regulating the timing and level of *cusC* induction in response to the environmental copper condition, and the acquirement of *cueO* makes it possible to safeguard against Cu(I) toxicity and to incorporate copper, resulting in rapid growth.

It has been reported that there are diversified systems for copper homeostasis in gamma proteobacteria (Hernández-Montes et al., 2012). Particularly, the periplasmic proteins working with CopA and CusC are diversified, probably to adapt to the different environments, during the evolution of gamma proteobacteria. Multicopper oxidases are encoded in the genomes of other Gram-negative bacteria. For example, *Salmonella enterica* sv. Typhimurium has a homologous CueO (alias CuiD), which demonstrates 80% identity over 516 amino acids with CueO (Osman and Cavet, 2011). Similar to the biochemical activity of CueO, CuiD has high oxidase activity for Cu(I), and reduces the concentration of periplasmic Cu(I) to avoid its toxic effects (Osman et al., 2010). The *cus* system is not present in *S. enterica* sv. Typhimurium (Pontel and Soncini, 2009). It is conceivable that instead of the *cus* system, another periplasmic protein CueP is involved in copper sequestration under anaerobic conditions, and constitutes a major cellular copper pool (Pontel and Soncini, 2009; Dupont et al. 2011). Although the mechanism by which CueP contributes to copper tolerance has not been fully understood, the CueP-like proteins are found in other Gram-negative bacteria such as *Yersinia* sp., *Citrobacter* sp., *Erwinia* sp., and *Corynebacterium* sp. (Osman and Cavet, 2011), indicating that most Gram-negative bacteria may have a multicopper oxidase and other periplasmic protains participated in the copper homeostasis.

Thus, Gram-negative bacteria could acquire periplasmic proteins worked with CopA and CusC to protect the cells against Cu(I) toxicity. In particular, the acquirement of multicopper oxidase may help to utilize the periplasm as a sensor and store for copper, resulting in expansion of living niche with a wide range of environmental copper concentrations.

## Materials and Methods

### Strains and Culture Condition

The *E. coli* K-12, MG1655 and MC4100 strains were used as the parental strains, and all strains and plasmids used in this study are presented in Tables S3 and S4, respectively. Primers used for genetic manipulations are presented in Table S2. The gene disruptions were performed with one-step inactivation method (Datsenko and Wanner, 2000). BW25113 pKD46 was used to construct Δ*cueR*::*cm*, Δ*cueR*::*km*, Δ*copA*::*cm*, Δ*copA*::*km*, Δ*cueO*::*cm*, Δ*cueO*::*km*, Δ*cusCFBA*::*cm* and Δ*cusCFBA*::*km* by transformation using PCR products amplified with the template plasmids, pKD3 and pKD4 and the primers, copA-del-f, copA-del-r, cueO-del-f, cueO-del-r, cusCFBA-del-f and cusCFBA-del-r. The Δ*cusRS*::*km* strain has already been constructed by the authors (Oshima et al., 2002). The strains expressing 3x Flag-tagged CusR and CueR were constructed by the epitope tagging procedure (Uzzau et al., 2001). BW25113 pKD46 was transformed with the DNA fragments encoding 3x Flag followed by the kanamycin resistant gene amplified with the template plasmid, pSUB11, and the primers, CueR-3xFLAG, CueR-RV, CusR-3xFLAG and CusR-RV. The derivatives of MG1655 and MC4100 were constructed by P1 transduction using P1 lysates prepared from the BW25113 derivatives disrupted genes or expressed Flag-tagged genes. The strains for the β-galactosidase analysis of the *cusR* promoter activity were constructed by the transduction of the lambda phage carrying the *cusR-lacZ* gene fusion into MC4100. The lambda phage was prepared by *in vivo* transfer of the DNA segment from the pMC1403 plasmid carrying the DNA fragment from 182 bp upstream to 27 bp downstream (corresponding to N-terminal 9 amino acids of CusR protein including the first Met) from the first adenine in the *cusR* gene, which expressed CusR (9 amino acids) – LacZ fusion protein, to *λRZ*5 phage (Carter-Muenchau and Wolf, 1989) by the RecA-mediated recombination as described previously (Ito et al., 1993). The DNA fragment to make the *cusR-lacZ* gene fusion was amplified by PCR with the *E. coli* chromosome DNA and the primers, ylcA3 and ylcA5, and then cloned into pMC1403 as *BamH*I digested fragment. The strains used for the Lux assay were prepared by the transformation of MG1655 and its derivatives with the luciferase reporter plasmids, constructed from pLUX. The DNA fragments including the promoters of *copA, cueO* and *cusC* were amplified by PCR using the primers, copA-promoter-01, copA-promoter-02, cueO-promoter-01, cueO-promoter-02, cusC-promoter-01 and cusC-promoter-02, and the *E. coli* chromosome DNA, and then cloned into pLUX as *Xho*I/*BamH*I fragments.

Luria-Bertani broth was used for the strain constructions. ChIP-chip and Lux assay were performed using Defined Medium A (DMA: 11.3 g K2PO4, 5.4 g NaH2PO4, 200 mg MgSO4, 10 mg CaCl2, 5 mg FeSO4, 0.5 mg ZnSO4, 0.5 mg MnSO4, 0.1 mg CuSO4, 0.1 mg CoCl2, 0.1 mg sodium borate, 0.1 mg sodium molybdate, 0.26 g EDTA, 2 g NH4Cl per litre) (Pirt, 1967). DMA was supplemented with 0.4% (w/v) glucose, and CuSO4. CuSO4 was buffered at pH 7 using 4 M glycine to avoid the precipitation of Cu(II) salts (Kershaw et al., 2005). MacConkey agar plate (BD Difco, Franklin Lakes, NJ) was used to monitor the *cusR* expression.

### Lux Assay

The luciferase (Lux) assay was performed according to the method used in our previous analysis of the response to zinc (Takahashi et al., 2015). MG1655 (a parental strain) and its derivatives, Δ*copA*::*cm* (SK14), Δ*cusCFBA*::*cm* (SK16), and Δ*cueO*::*cm* (SK18) (Table S3) were transformed with the plasmids, pLUX, pLUX*copA*, pLUX*cueO* and pLUX*cusC*. The strains were pre-cultivated in 1 mL of DMA with 0.4% glucose, 25 µg/mL kanamycin, and 8 mM glycine at 37°C overnight. The strains were then diluted to the OD600 value of 0.04 in fresh 2 mL of DMA and re-cultivated for 2.5 h. After the cultures reached at early log phase (OD600 = 0.1∼0.4), 180 µL of each culture solution was transferred to Greiner 96-well microwell white plate (Greiner, Frickenhausen, Germany) and pre-incubated with shaking at 37°C for 15 min. Adequate amount of copper and glycine solution prewarmed at 37°C was added to each well at final concentrations from 0 µM (only glycine solution was added) to 400 µM of copper so that the total volume became 200 µL. Following addition of copper, the luciferase activities by absorbance (OD595) were measured every 30 seconds for one hour using a Mithras LB940 (Berthold Technologies, Bad Wildbad, Germany) (Table S5). To normalize the bioluminescent activity per unit cell mass, the luminescence activity was divided by the absorbance of cell culture.

### ChIP-chip Analysis

All strains used for ChIP-chip analysis were grown in 10 mL of DMA with aeration at 37°C until the culture reached at the OD600 value of 0.8. Each culture was fixed with formaldehyde at a final concentration of 1% for 20 min at 37°C. The excess formaldehyde was quenched with 1.5 mL of 3 M glycine for 5 min. Cultures were harvested and subjected to successive washing steps with TBS and lysis buffer (10 mM Tris-HCl pH 8.0, 20% sucrose, 50 mM NaCl, 10 mM EDTA). Cells were suspended in 500 µL of lysis buffer containing 20 mg/mL lysozyme and incubated for 30 min at 37°C and keep them at –30°C until the next step. 4 mL of IP buffer (50 mM HEPES-KOH pH 7.5, 300 mM NaCl, 1 mM EDTA, 1% Triton X-100, 0.1% sodium deoxycholate, 0.1% SDS, 5% glycerol) and 50 µL of 100 mM PMSF were added to the samples. The samples were then sonicated by a XL2020 (Astrason, Plainview, NY) on ice for 2 min (4 x 30 s with 30 s intervals) to prepare CueR samples or 10 min (20 x 30 s with 30 s intervals) to prepare CusR samples. To prepare the supernatant (whole cell extract) including chromatin, cell samples was clarified by centrifugation at 20,000 g for 30 min at 4°C. The supernatant was mixed with 15 µL of protein A Dynal Dynabeads 100.02 (Invitrogen, Waltham, MA) coated with anti-Flag antibody; to coat the protein A beads with antibody, the beads were washed twice with 500 µL of cold TBS containing 5 mg/mL BSA, and coated with anti-FLAG M2 antibody (Sigma-Aldrich, St. Louis, MO) by gentle mixing with 20 µL of cold TBS containing 5 mg/mL BSA added 1.1 µL of antibody solution (4.6 µg/µL) for 6 h at 4°C before use. The mixture was further incubated at 4°C overnight with rotation. The beads were rinsed once with 1.5 mL of IP buffer, once with 1 mL of IP buffer for 5 min with rotation at 4°C, once with 1 mL of IP salt buffer (IP buffer containing 500 mM NaCl), three times with 1 mL of wash buffer (10 mM Tris-HCl pH8.0, 250 mM LiCl, 1 mM EDTA, 0.5% Nonidet P-40, 0.5% sodium deoxycholate) and then once with 1 mL of TE (10 mM Tris-HCl, 1mM EDTA pH 8.0). After the rinses, the protein-DNA complexes bound on the beads were eluted by 100 µL of elution buffer (250 mM Tris-HCl pH 7.5, 50 mM EDTA pH 8.0, 5% SDS). DNA fragments crosslinked with proteins were decrosslinked by heating at 65°C for 20 min to prepare purified DNA fragments from whole cell extract and immunoprecipitated fractions. After decrosslinking, proteins in whole cell extract and immunoprecipitated fractions were digested with 2 mg/mL proteinase K (Takara Bio Inc., Kusatsu, Japan) at 42°C for 2 h, followed by incubation at 65°C for 6 h to inactivate proteinase K. Free DNA fragments in whole cell extract and immunoprecipitated fractions were purified by the QIAquick PCR Purification Kit (QIAGEN, Hilden, Germany) and eluted with 50 µL of elution buffer provided by the kit.

Random amplification of purified DNA fragments was performed according to Katou et al. (2006). Terminal labeling of amplified DNA purified from whole cell extract and immunoprecipitated fractions, and hybridization to the microarray were performed according to the Affymetrix instruction (Affymetrix, Santa Clara, CA). The custom Affymetrix oligonucleotide array (ECTILE2b520514F) was employed to obtain raw data (CEL.file). Raw data was visualized by In Silico Molecular Cloning program array edition (In Silico Biology, Yokohama, Japan) to identify the protein binding regions. Signal intensities of mismatch probes were subtracted from those of perfect match probes. Probes with a negative value for signal intensity were excluded from further analysis. The mean values of signal intensities of DNA isolated from immunoprecipitated fraction and whole cell extract fraction without purification (control DNA) were normalized to 500. Then, the signal intensities of DNA in the immunoprecipitated fraction were divided by those of control DNA. In order not to use the abnormal values caused by the low signal intensities of control DNA signal intensities in the analysis, we filtered 10% of probes with lower intensities of control DNA samples by the global rate trimming function in In Silico Molecular Cloning program. As we found the high background noise likely due to the PCR amplification (especially CusR data), the binding sites were identified by eye inspection.

### Growth Measurements

Optical density measurements were taken from each strain type in each media type in triplicate using a SmartSpec^TM^ Plus spectrophotometer (Bio-Rad Laboratories, Inc., Hercules, CA). 2 µL of stationary-phase culture grown in DMA was added to 2 mL of DMA with final concentrations of either 0 µM, 50 µM, 100 µM, 250 µM, 500 µM, 750 µM, 1,000 µM, 2,000 µM, 3,000 µM, 4,000 µM, 5,000 µM, or 6,000 µM in a 15 mL sterile plastic tube. The cultures were grown for 8 h at 37°C with shaking at 185 rpm.

### Mathematical Model

We developed a mathematical model that describes the molecular process of copper homeostasis representing the dynamics of Cu(I) concentration in the cytoplasm and the periplasm (Figure 1A). There are 16 variables; the concentration of CopA, the concentration of CueO, the concentration of CusC, the concentration of active (copper-bound) CueR (active CueR), the concentration of CusS, the concentration of active (phosphorylated) CusS (CusS-P), the concentration of free (ionic) Cu(I) in the cytoplasm (Cuc^+^), the concentration of free Cu(I) in the periplasm (Cup^+^), the concentration of free Cu(II) in the periplasm (Cup^2+^), the concentration of copper bound to the reservoir molecules, i.e. any other copper-binding molecules in the periplasm (*R*), the concentration of CusR, the concentration of phosphorylated CusR (CusR-P), the concentration of phosphorylated CusR dimer (CusR-P dimer), the concentration of protein complex of phosphorylated CusS and CusR (CusS-P·CusR), the concentration of protein complex of CusR and phosphorylated CusR (CusS·CusR-P), and the concentration of active (copper-bound) phosphorylated CusS (CusS-2Cu_p_^+^-P). The equations are

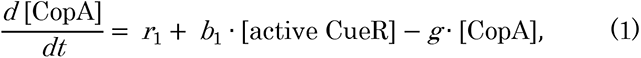

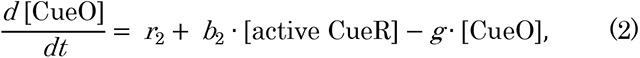

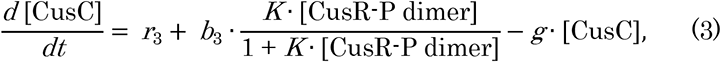

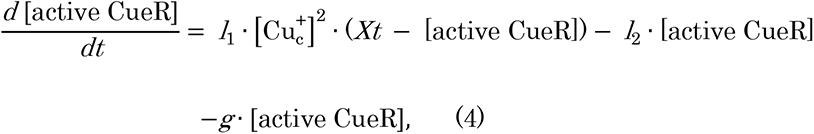

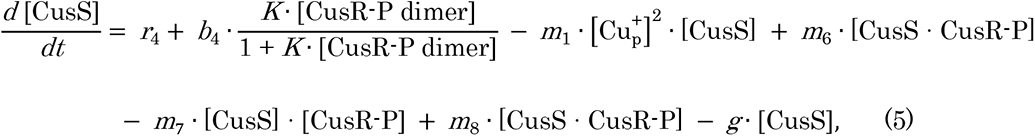

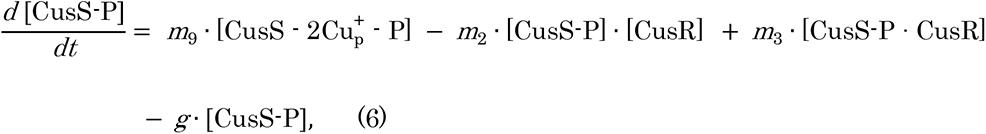

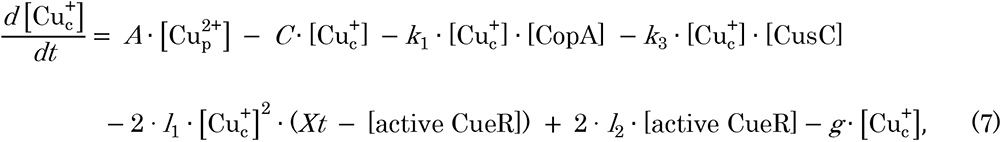

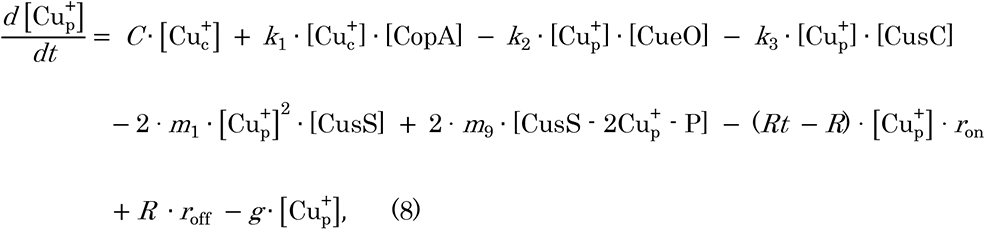

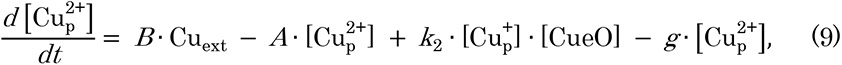

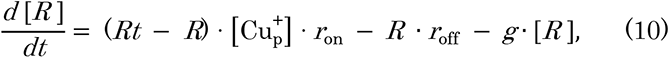

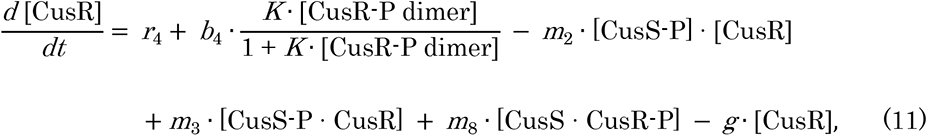

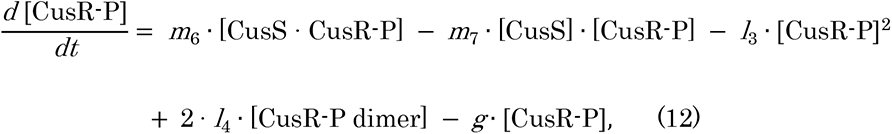

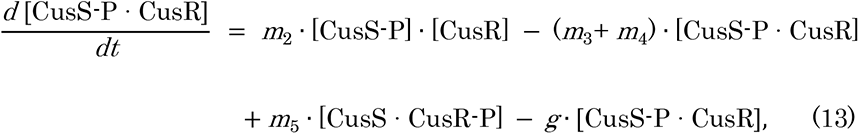

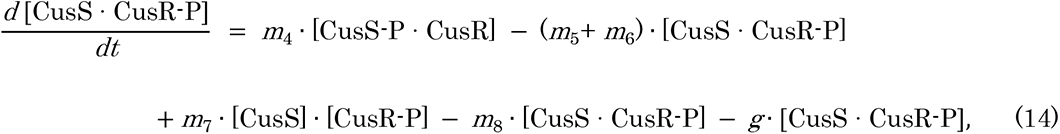

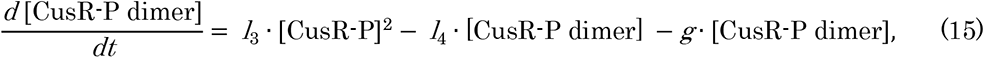

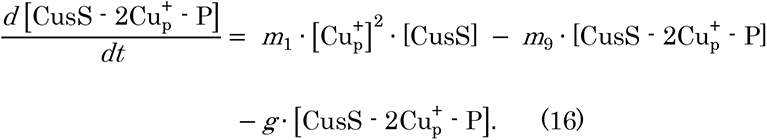

CopA is produced at a basal rate *r_1_* and activated by active CueR with a constant of proportionality *b_1_* (equation 1). CopA and all other modeled components are diluted due to cell growth at rate *g*; the four strains examined could potentially grow at different rates so the value of *g* is strain-dependent. CueO is produced at a basal rate *r_2_* and activated by active CueR with a constant of proportionality *b_2_* (equation 2). CusC is produced at a basal rate *r_3_* and activated by CueR-P dimer with a constant of proportionality *b_3_* and activation coefficient *K* (equation 3). *XT* represents the total amount of CueR (assumed to be constant), and so inactive CueR is given by *XT*−active CueR. The binding of two copper ions is required to convert inactive CueO to active CueO at rate *l_1_*. The active form can also revert to the inactive form at rate *l_2_* (equation 4). The kinetic parameters associated with the phosphorylation cycle model (Figure S2) are introduced (equations 5, 6, 11–16). Briefly, CusS is phosphorylated after binding to copper (equations 5, 6 and 16). CusR is phosphorylated by forming the complex with CusS-P (equations 11–14). CusR-P dimer activates the expression of CusC, CusS and CusR (equation 15). CusS is produced at a basal rate *r_4_* and activated by CusR-P dimer with a constant of proportionality *b_4_* (equation 5). The binding of two copper ions is required to form the copper-CusS complex (CusS-2Cu^+^p) at rate *m_1_* (equations 5 and 16), and two copper ions dissociates from the copper-CusS complex to phosphorylate CusS at rate *m_9_* (equations 6 and 16). The phosphorylated CusS binds to CusR at rate *m_2_* (equations 6, 11, and 13), and the protein complex can be dissociated at rate *m_3_* (equations 6, 11, and 13). The phosphorylated CusS phosphorylates CusR at rate *m_4_* (equations 13 and 14), and the irreversible reaction can be proceeded at rate *m_5_* (equations 13 and 14). The protein complex is dissociated at rate *m_6_* to produce the phosphorylated CusR (equations 5, 12, and 14). In this model, CusS also dephosphorylates CusR; CusS binds to the phosphorylated CusR at rate *m_7_* (equations 5, 12, and 14), and dephosphorylates at rate *m_8_* (equations 5, 11, and 14). In equation 15, phosphorylated CusR forms dimer at rate *l_3_*, and the dimer is dissociated at rate *l_4_*. In equations 7–9, the copper spatiotemporal dynamics in the cytoplasm and the periplasm is described. Cu(II) enters bacterial cells without any specific copper uptake system, and therefore, import of Cup^2+^ is proportional to the concentration of external copper with rate *B* (equation 9). The diffusion from the periplasm into the cytoplasm is proportional to the concentration of Cup^2+^ with rate *A*, and Cup^2+^ is generated by the oxidation of Cup^+^ by CueO with a constant probability *k_2_* (equations 8 and 9). The export of Cuc^+^ is proportional to the concentration of Cuc^+^ with a basal rate *C*, and linearly dependence on CopA with a constant of proportionality *k_1_* (equations 7 and 8). CusC exports Cup^+^ and Cuc^+^ into the outside of cells with a constant of proportionality *k_3_* (equations 7 and 8). In equation 10, the Cup^+^-unbound and Cup^+^-bound reservoir are described. *RT* represents the total amount of reservoir molecule in the periplasm (assumed to be constant), so Cup^+^-unbound reservoir molecule is given by *RT*−*R*. The reservoir molecule binds to Cup^+^ at rate *ron*, and the Cup^+^-bound reservoir can be dissociated at rate *roff*.

### Monte Carlo Simulations

Parameter estimations were carried out using the Metropolis-Hastings algorithm (Hastings, 1970; Herman et al., 2011). For the majority of parameters, uninformative priors have been used. As proposal distributions, a lognormal distribution was used and the variances of the distributions were empirically chosen in order to ensure acceptance probabilities close to 0.25–0.30. The parameters were updated separately in each step, and 10,000,000 iterations were carried out. ODE calculations were performed by the cvode solver with Newton iterations provided by the Sundials library (Cohen and Hindmarsh, 1996).

### Calculations of ODE and Stochastic Simulations

ODE calculations were performed by the deSolve package (Soetaert et al., 2010) using the statistical environment *R*. Stochastic simulations based on the Gibson-Bruck algorithm (Gibson and Bruck, 2000) were performed until 1.1 × 10^6^ s, i.e. 300 h, using Dizzy (ver. 1.11.3) (Ramsey et al., 2005).

### Comparative Genome and Phylogenetic Analyses

Orthologous groups of all genes in *E. coli* (GCF_000005845.2), *Synechococcus elongatus* (GCF_000010065.1), *Bacillus subtilis* (GCF_000009045.1) and *Staphylococcus aureus* (GCF_000013425.1) were identified by using OrthoFinder (ver. 2.3.3) with default parameters (Emms and Kelly, 2019). Phylogenetic analysis was conducted based on 16S rRNA sequences. All 16s rRNA sequences were aligned by using MAFFT (ver. 7) with default parameters (Katoh et al., 2019). The phylogenetic tree was constructed by the neighbor-joining method implemented the MEGA software (ver. 10.1.5) with the Kimura two-parameter model (Kimura et al., 1980; Kumar et al., 2018). Bootstrap values were calculated with 100 resamples.

### Calculation of *R*^2^ value

We calculated the Pearson correlation coefficient between the experimental and theoretical values (log_2_ ratio) after addition of copper until 60 min.

### Calculation of Coefficient of Variation, and Lower/Upper Bounds of Each Parameter

All model parameters were estimated from the posterior distributions of the MCMC runs. Table S1 shows the median value for each parameter, the coefficient of variation, and lower and upper bounds, that have been calculated by subtracting or adding 1.96 times the standard deviation from the median log value. This process has been carried out to avoid sampling errors at the extreme values of the Markov chain, assuming a Gaussian posterior centered on the median value.

## Acknowledgements

We thank Dov Stekel and Jon Hobman for critically reading the manuscript. We also thank Masatoshi Kawabata and Berthold Japan for supporting Lux assay using a Mithras LB940. This work was partly supported by the Japan Science and Technology Strategic Japanese-UK Cooperative Program on Systems Biology to N.O., T.O. and H.T., MEXT KAKENHI (24780072 and 16K18671), AMED under Grant Number JP19fm0208024 to H.T., MEXT KAKENHI (16H06279) to H.T. and J.I., and MEXT KAKENHI (17K15151) to J.I.

## Authors’ Contributions

J.I., T.O. and H.T. conceived the project. All members performed the experiments and analyzed the experimental data. J.I., T.O. and H.T. developed the model and performed the numerical simulation. J.I., T.O. and H.T. wrote the manuscript. TM, CK and SK equally contributed to wet works, including construction of most of strains used in this study, ChIP-chip, *lacZ* and Lux reporter assays. S.I. performed ChIP-chip experiments with T.M. H.A. constructed Flag tagged strains. T.O., S.I. and N.O. initiated the project.

---

**Figure S1.**
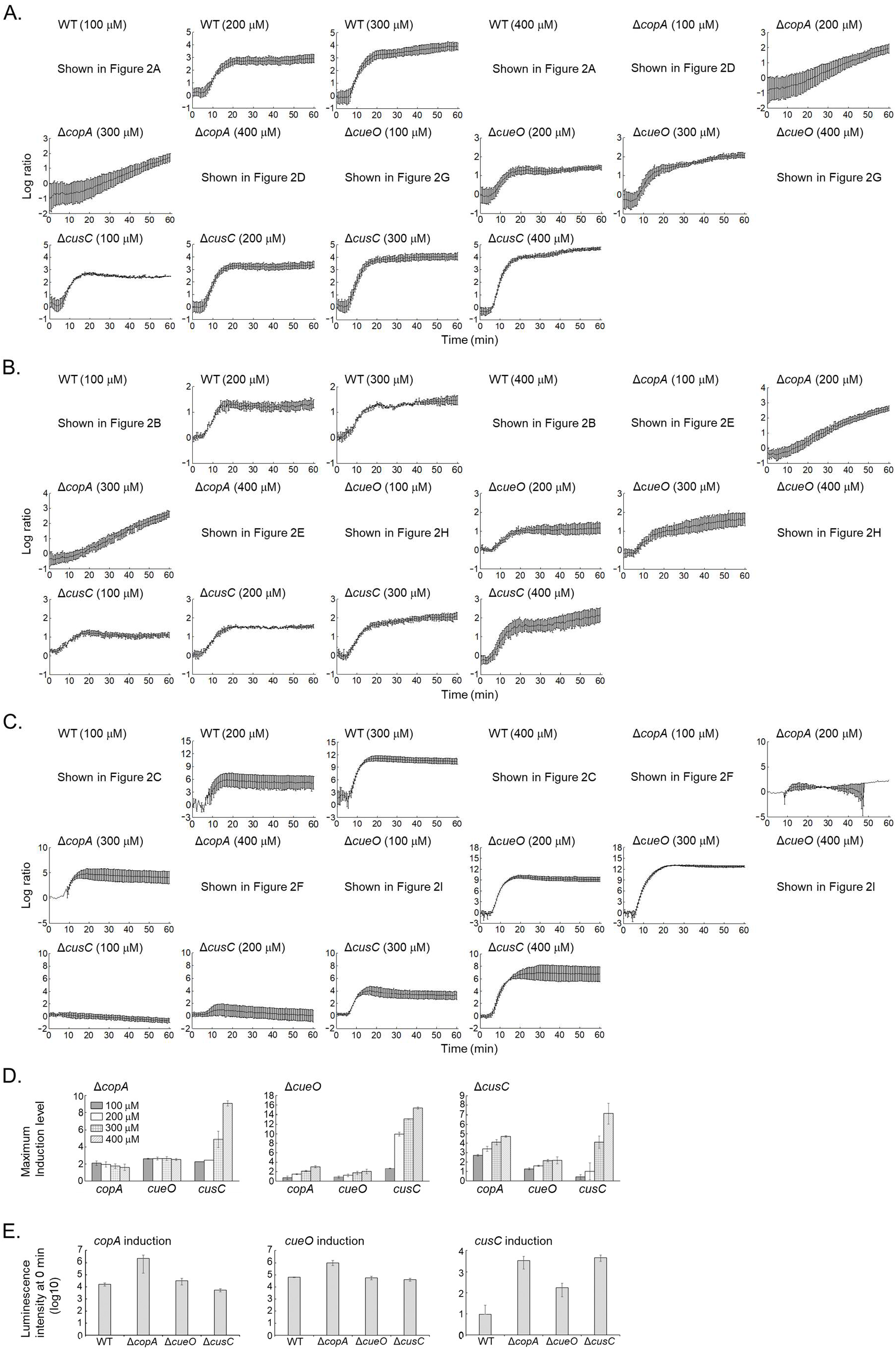
(A–C) Time series of induction of *copA*, *cueO*, and *cusC* promoters following addition of copper, log to base 2 ratios normalized to the value at *t*=0. Error bars indicate s.d. (*n*=3). (D) Maximum induction levels of P*copA*, P*cueO*, and P*cusC* relative to the data at 0 min in the Δ*copA*, Δ*cueO*, and Δ*cusC* strains (log to base 2 ratios to the value at *t*=0). (E) Luminescence intensities of *copA*, *cueO* and *cusC* promoters in the WT, Δ*copA*, Δ*cueO*, and Δ*cusC* strains at *t*=0. The luminescence intensities correspond to the data in DMA^400^. Error bars indicate s.d. (*n*=3).

**Figure S2.**
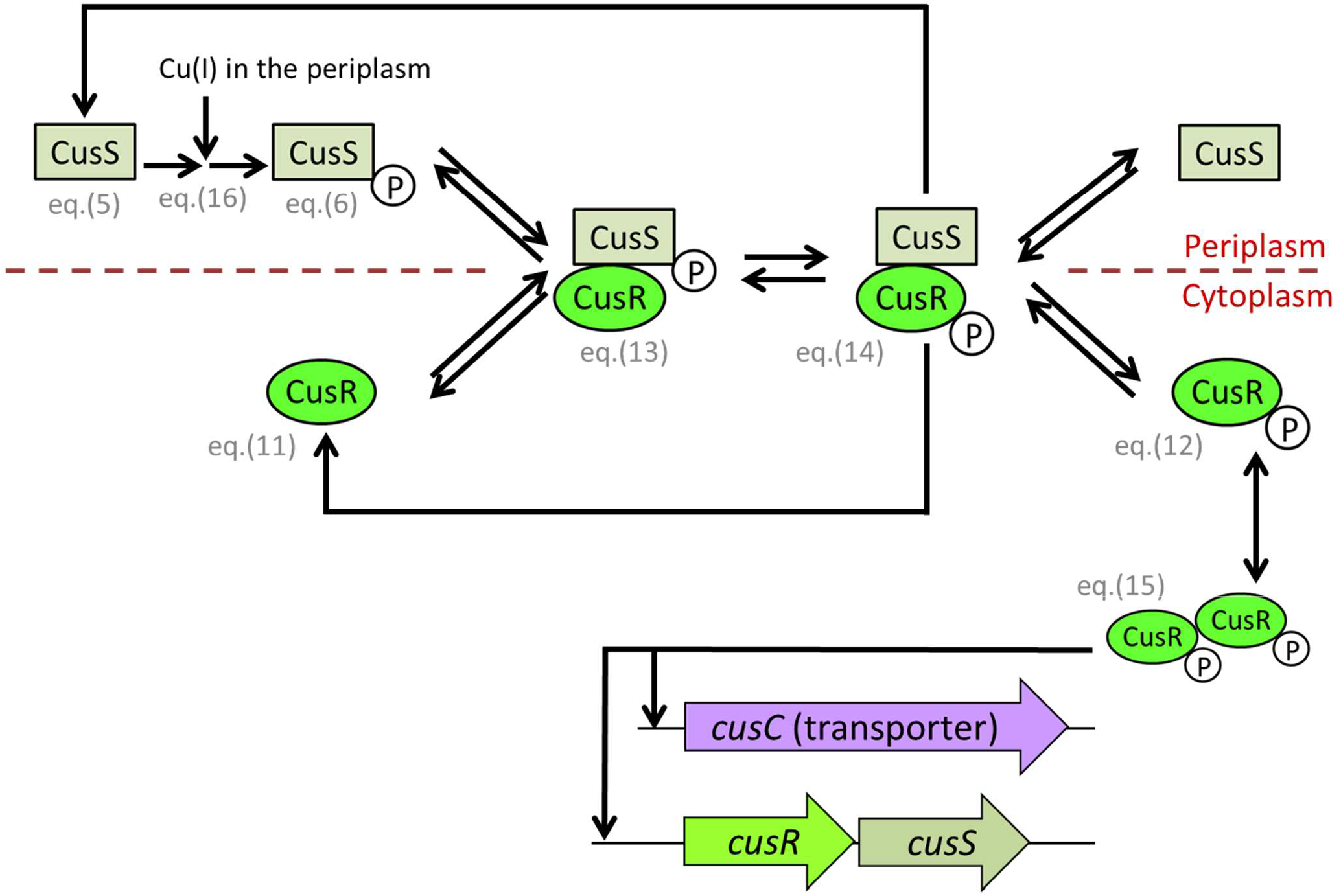
A schematic figure of *cusRS* regulation. The phosphorylation and dephosphorylation ability of CusS, and the positive feedback by the phosphorylated CusR are indicated by arrows. P denotes being phosphorylated. The reaction scheme includes the association, dissociation, phosphorylation and dephosphorylation.

**Figure S3.**
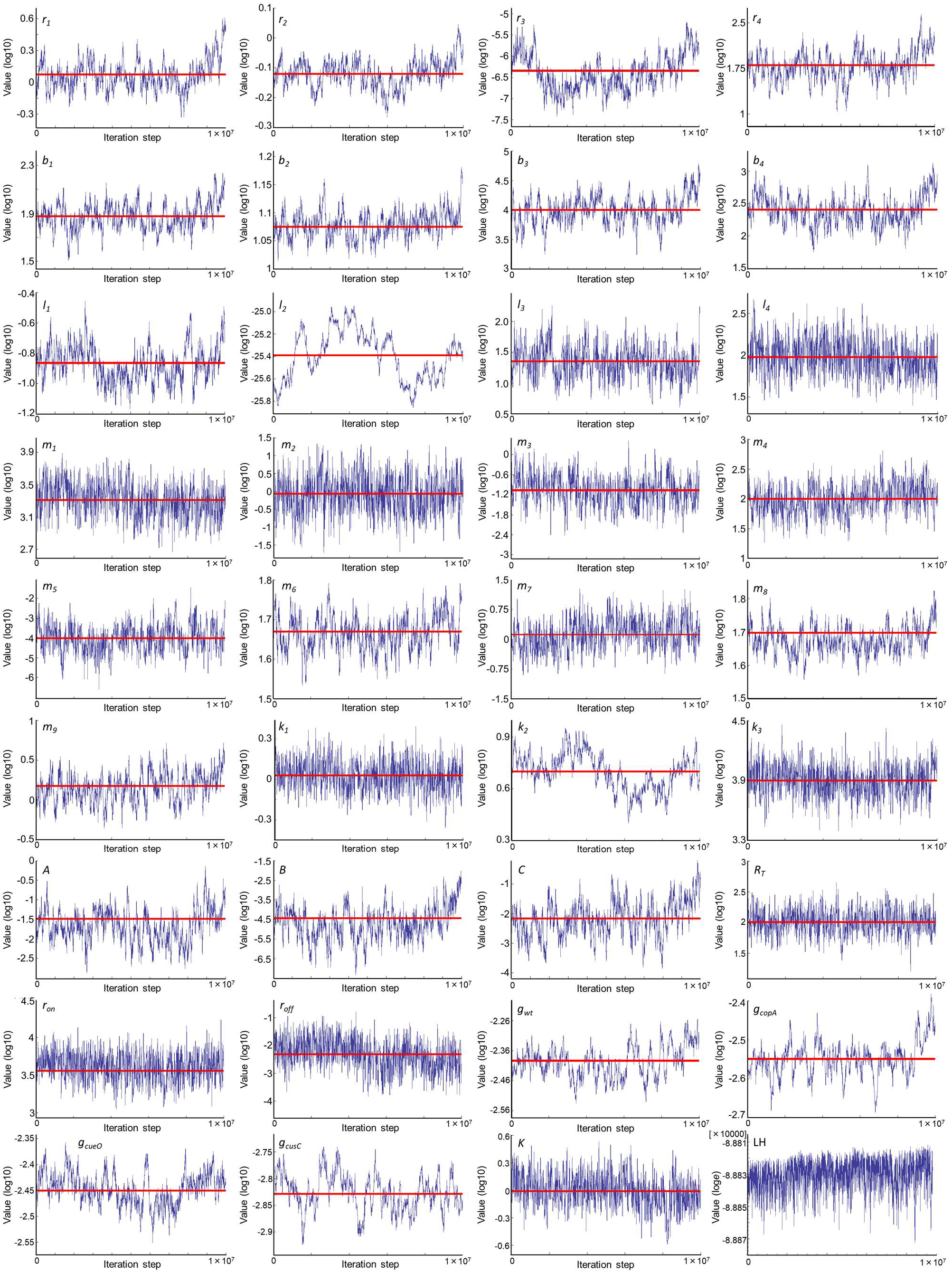
Trace plots for MCMC runs. 35 parameter (log to base 10) and natural logarithm likelihood were plotted. Blue line is the median value of posterior distribution.

**Figure S4.**
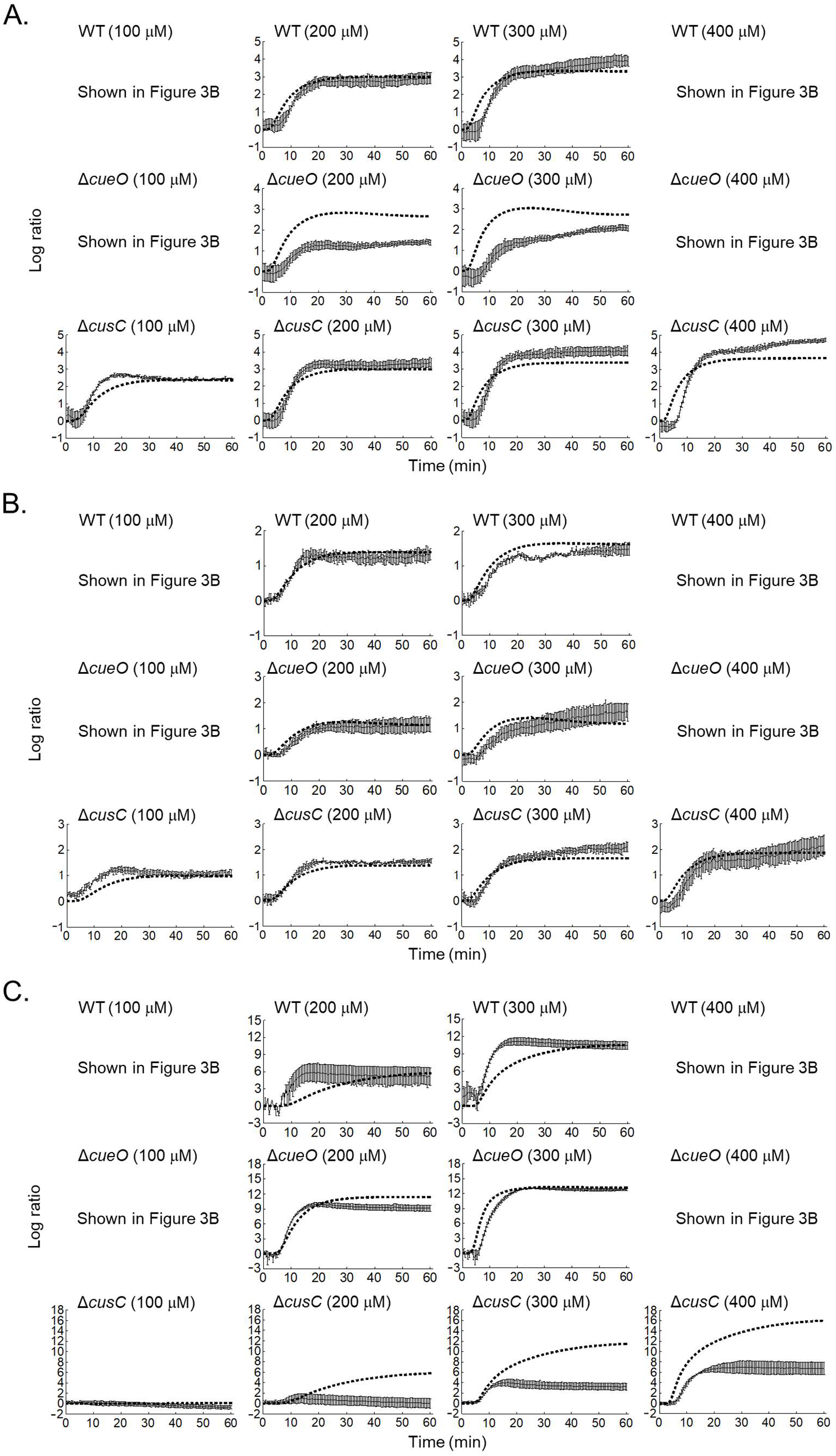
Time series of induction of (A) *copA*, (B) *cueO*, and (C) *cusC* promoters following addition of copper, log to base 2 ratios normalized to the value at *t*=0. The sold line indicates the experimental data (error bar is s.d.), which are already shown in Figure S1. The dotted line indicates the theoretical data with the best fit parameters.

**Figure S5.**
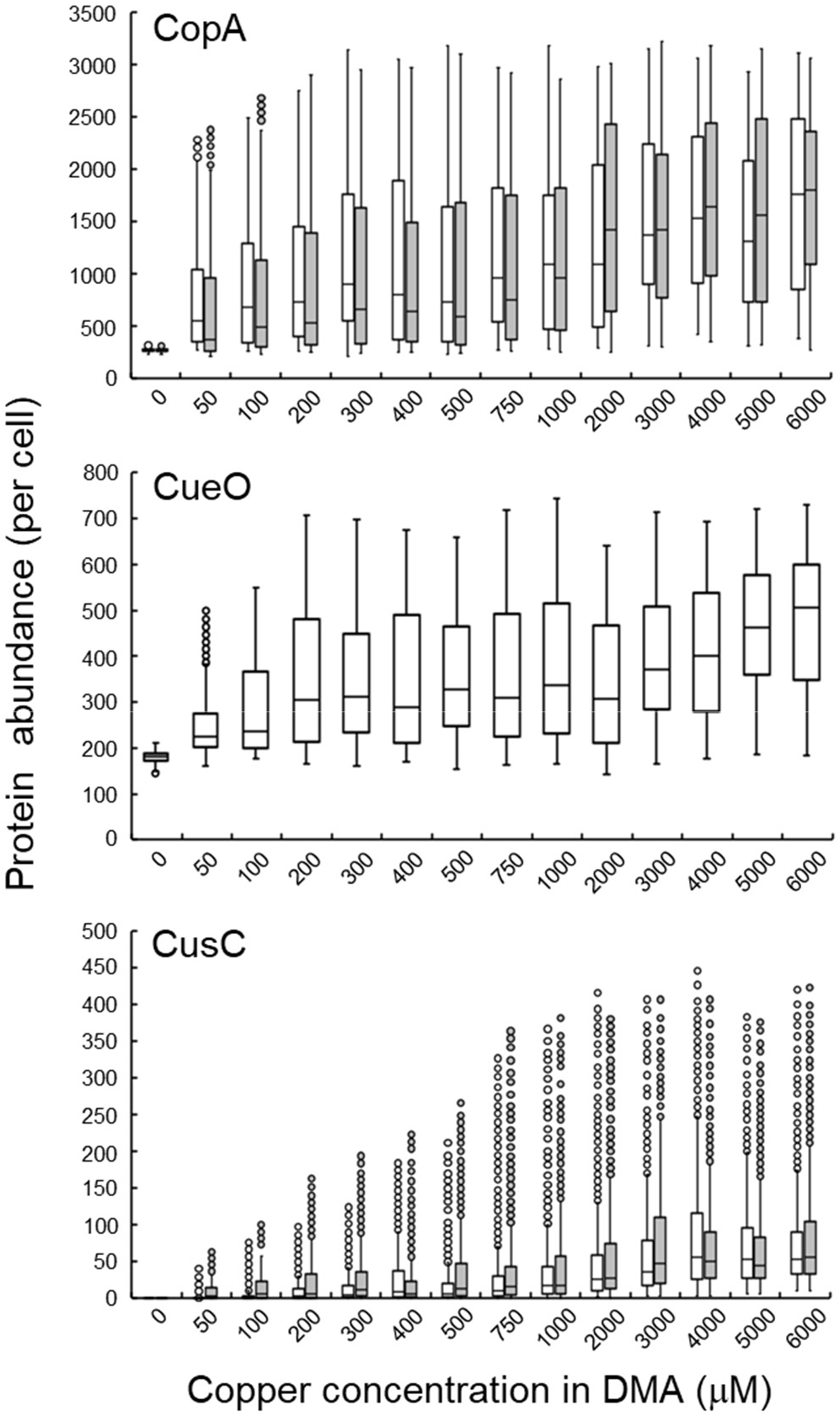
Boxplots of abundance of CopA, CueO, and CusC in the WT (black) and Δ*cueO* (gray) strains.

**Figure S6.**
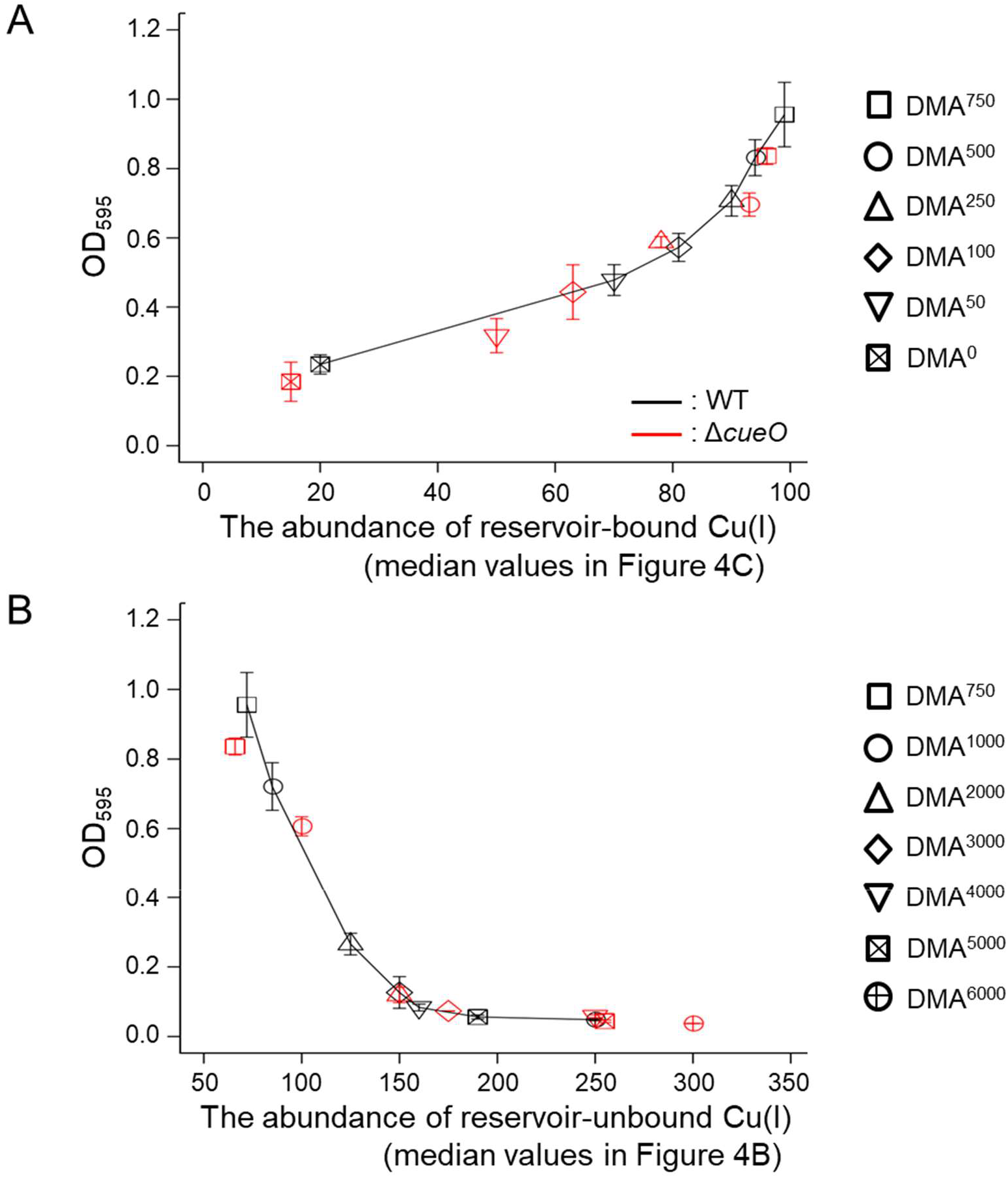
(A) Relationship between the growth and the abundance of reservoir-bound copper in the WT (black) and Δ*cueO* strains (red) in DMA^0^–DMA^750^. The vertical and horizontal axes correspond to the OD595 values, and the median value of abundance of reservoir-bound copper, respectively. (B) Relationship between the growth and the abundance of periplasmic free copper in the WT (black) and Δ*cueO* strains (red) in DMA^750^–DMA^6000^. The vertical and horizontal axes correspond to the OD595 values, and the median value of periplasmic free copper.

**Table S1.** Estimates of all model parametersT

**Table S2**. A list of primers used in this study

**Table S3**. A list of bacterial strains used in this study

**Table S4.** A list of plasmids used in this study

**Table S5.** Promoter activities of *copA*, *cueO*, and *cusC* in the WT, Δ*copA*, Δ*cueO*, and Δ*cusC* strains measured by the Lux reporter system

